# Single-cell epigenomic tracing of lifelong endothelial cell plasticity across mouse organs

**DOI:** 10.1101/2021.05.12.443777

**Authors:** Xianhong Yu, Yaxi Liu, Xiaoge Liu, Haiqing Xiong, Aibin He

**Affiliations:** Beijing Key Laboratory of Cardiometabolic Molecular Medicine, Institute of Molecular Medicine, College of Future Technology and Peking-Tsinghua Center for Life Sciences, Peking University, Beijing 100871, China

## Abstract

Endothelial cells (ECs) across ages and tissues are highly heterogeneous in developmental origins, structures, functions, and cellular plasticity. Here, we applied CoBATCH for single-cell epigenomic tracing of dynamic EC lineage histories in five mouse organs from development to ageing. Our analyses showed that epigenomic memory reflects both developmental origins and tissue-restricted specialization of EC sublineages but with varying time lengths across organs. To gain insights into cellular plasticity of ECs, we identified bivalent chromatin occupancy of otherwise mutually exclusive EC- (ERG) and mesenchymal-specific (TWIST1/SNAI1) transcription factors promoting endothelial-to-mesenchymal transition. We further revealed that pseudotime trajectories by histone modifications H3K36me3 and H3K27ac faithfully recapitulate short- and long-range EC fate change over senescence, respectively. Together, our data provide a unique exploration of chromatin-level cell fate regulation of organotypic EC lineages across the lifespan.

**One-Sentence Summary:** Single-cell chromatin binding is examined for tracing endothelial cell lineages in mouse organs across the lifespan.

---

Vascular endothelial cells (ECs) emerge de novo from both the extraembryonic yolk sac and mesodermal progenitors during early embryo development, and further undergo specification and remodeling to create distinct vessel subtypes required for tissue development, homeostasis, and regeneration(*1-8*). Importantly, considerable EC heterogeneities for structure, phenotype and function manifest across tissues from development to ageing(*9-13*). Single-cell transcriptomic studies have largely improved our understanding of EC heterogeneities in mouse organs at the certain stages(*14-20*), offering an unprecedented opportunity in cataloging tissue-specific EC subtypes. ECs from different tissues exhibit high expression of respective tissue-restricted transcription factors, such as *Foxq1* in brain, *Hoxd9* in testis, *Foxf1* in lung, *Pparg* in skeletal muscle, and *Gata4* in liver(*15*). Recently, single-cell epigenomic approaches, such as chromatin accessibility by scATAC-seq (assay for transposase-accessible chromatin with high-throughput sequencing), have begun to reveal regulatory heterogeneities during cellular diversification in tissues(*21-24*). Emerging significant questions include whether and how these tissue-restricted features are perpetuated in the epigenome during development or across the lifespan. Whether tissue microenvironment educates ECs to support the parenchymal cells and their functions or simply they share the common progenitors to some extent recently attracts intensive investigation(*25*). Here, we applied our recently developed CoBATCH(*26*) for measuring single-cell chromatin binding of histone modifications and transcription factors as a surrogate for tracing the EC lineage dynamics across mouse organs (brain, heart, lung, kidney and liver) from embryonic day E16.5 to ageing. Our data present a comprehensive epigenomic landscape for deciphering developmental origin and specialization of EC phenotypes and cooperative TF actions in regulation of cell fate decision at the single-cell level.

## Single-cell CoBATCH profiling of EC lineages in five mouse organs across development and ageing

With an improved CoBATCH protocol(*26*) (See also Materials and methods), we first performed single-cell profiles of endothelial cell (EC) lineages labeled by an EC-specific marker gene *Cdh5* (encoding VE-Cadherin)(*27*) from five organs (heart, brain, liver, lung, and kidney) at the stages of embryonic day (E)16.5, neonate (P5), adulthood (3-4 months), and senescence (17-19 months) (table S1). Tissues from 4-day tamoxifen-induced Cdh5^CreERT2^::Rosa26^tdTomato/+^ mice were dissected and dissociated before FACS (fluorescence-activated cell sorter) sorting for tdTomato^+^ cells. Thus, ECs and their derivatives were subjected to single-cell CoBATCH profiling for H3K27ac, a histone mark for active enhancer (*28*) (Fig. 1A, table S2 and S3). The EC labeling was verified by immunostaining for CD31 (fig. S1, A). Together, we obtained H3K27ac profiles of 10,527 single cells passing quality filter across the developmental and aged stages, yielding 6,255 ± 2,568 deduplicated fragments (mean ± s.d.) per cell and a high fraction of reads in peaks (∼80%) for each sample (fig. S1, B to D, and table S4). Expectedly, strong H3K27ac signals in the vicinity of *Cdh5* and developmental gene *Gata4* were observed in hearts across all stages (fig. S2A). The data quality was also validated by the peak annotation and transcription factor (TF) motif enrichment. The peaks were mostly found at cis-regulatory regions including promoters and distal enhancers, and enriched for motifs of MEF2 and ETS families(*29*) (fig. S2, B and C). Thus, these CoBATCH data can be used to reconstruct a comprehensive epigenomic landscape while tracing dynamic EC lineages across organs with time.

**Fig 1.**
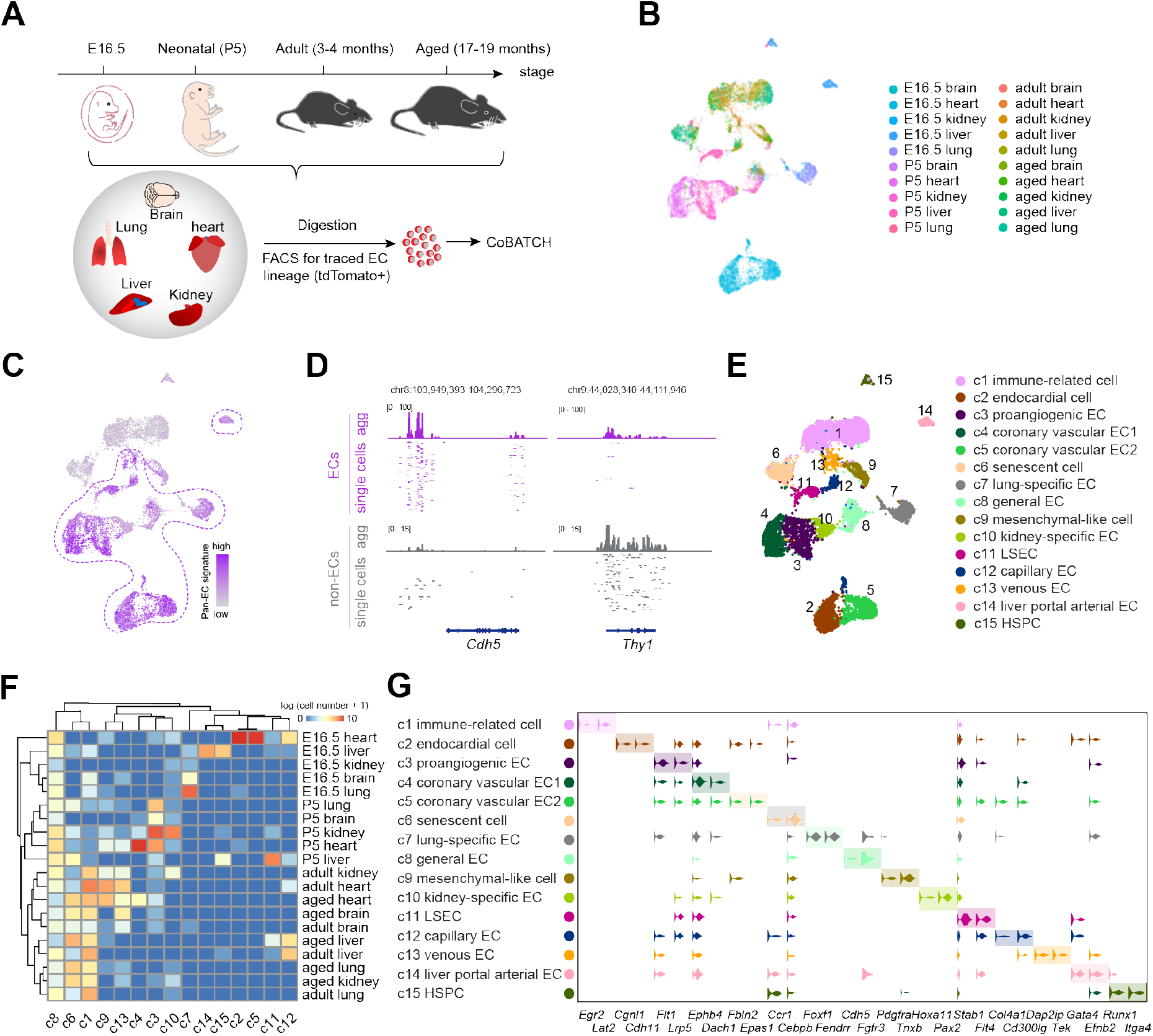
Single-cell CoBATCH profiling for epigenomic tracing of endothelial lineages across mouse organ development and ageing. **(A)** Scheme of experimental design. Cdh5^creERT2^::Rosa26^tdTomato/+^mice were administered with 2-dose tamoxifen for 4 days before sacrificed. Heart, brain, kidney, liver, and lung were microdissected and dissociated before FACS sorting for tdTomato^+^ cells for single-cell H3K27ac CoBATCH experiments. **(B)** UMAP visualization of H3K27ac profiling of 10,527 single cells traced by Cdh5^creERT2^::Rosa26^tdTomato/+^ from indicated organs at four stages. **(C)** Projection of pan-EC signature score on UMAP in (B). ECs and non-ECs were readily distinguished based on the signature score. ECs circled by dotted line presented the high pan-EC signature score. The pan-EC signature score was calculated using normalized H3K27ac signals in 137 peaks overlapped with the TSS ± 5 kb regions of 42 pan-EC signature genes (See also Methods). **(D)** Track view showing H3K27ac signals of aggregate and selected single ECs or non-ECs at representative EC marker gene *Cdh5* and mesenchymal marker gene *Thy1*. Purple color indicates ECs and grey color indicates non-ECs. Randomly selected 300 single cells are shown here in the descending order of signals, agg, aggregate. **(E)** UMAP visualization of the annotated clusters of all traced lineage cells from five organs at four stages. The clusters were identified by LSI clustering based on 57,488 H3K27ac peaks called from aggregate profiles. LSEC, liver sinusoidal endothelial cells; HSPC, haematopoietic stem and progenitor cells. **(F)** Heatmap showing hierarchical clustering of cell number (log2-transformed) in each cluster (in column) and each sample (in row). The color from blue to red represents the cell number from low to high. The clusters are annotated as in (E). **(G)** Violin plots showing the normalized H3K27ac signals at representative cell-type specific/enriched gene loci.

We next performed unsupervised dimensionality reduction using latent semantic indexing (LSI) algorithm followed by non-linear Uniform Manifold Approximation and Projection (UMAP) embedding to visualize single cells(*30*). Clustering of all EC lineage cells across tissues and ages presented prominent epigenomic heterogeneities (Fig. 1B). Two major populations were readily distinguished, corresponding to ECs and non-ECs, as demonstrated by projecting H3K27ac signals of pan-EC signature genes (Fig. 1C). Both the aggregate and single-cell profiles exhibited pronounced H3K27ac signals at the representative marker gene loci (*Cdh5* for ECs and *Thy1* for mesenchymal cells (MCs)(*31*)) (Fig. 1D). To further explore the heterogeneity of EC lineage cells across organs from development to ageing, 15 distinct subpopulations were identified and annotated by the H3K27ac signals around known marker genes as well as sampling information (organ and stage). Among 15 clusters, several were sourced from a single organ, such as endocardial cells (*Cgnl1*(*32*), *Cdh11(33*)), coronary vascular ECs (*Ephb4*(*34*), *Dach1*(*35*), *Fbln2(36), Epas1(37*)), lung-specific ECs (*Fendrr*(*38*), *Foxf1*(*39*)), liver portal arterial ECs (*Gata4*(*40*), *Efnb2*(*41*)), kidney-specific ECs (*Hoxa11*(*42*), *Pax2*(*43*)), and liver sinusoidal ECs (LSEC) (*Stab1*(*44*), *Flt4*(*45*)). Differing from the ECs restricted to particular tissues, functional proangiogenic ECs (*Flt1(46), Lrp5*(*47*)) were found in multiple organs. Similarly, c8 as an independent cluster comprising ECs from multiple organs across all stages, was thus defined as general ECs (*Cdh5*(*27*), *Fgfr3*(*48*)). Our data also separated ECs by vascular beds, such as capillary ECs (*Cd300lg(15), Col4a1(49*)) and venous ECs (*Tek*(*41*), *Dab2ip(50*)). Most immune-related cells (*Egr2*(*51*), *Lat2*(*52*)) and senescent cells (*Ccr1*(*53*), *Cebpb*(*54*)) were present at adult and aged stages, indicating a unique epigenetic role in mounting the immune response and ageing process(*14, 55*). Interestingly, we identified hematopoietic stem and progenitor cells (HSPC) (*Itga4*(*56*), *Runx1*(*57*)) in liver and mesenchymal-like cells (*Pdgfra*(*58*), *Tnxb*(*59*)), consistent with the expected cellular processes of the endothelial-to-hematopoietic transition (EHT) and endothelial-to-mesenchymal transition (EndMT) during those developmental stages (Fig. 1, E to G).

### Epigenomic memory reveals developmental origins and specialization of ECs

We asked whether single-cell trajectories established from 4-day genetic lineage tracing across all stages are capable to establish the link between the epigenomic memory of dynamic EC lineages and their developmental origin. We used UMAP to separately visualize single-cell H3K27ac CoBATCH profiles at the E16.5, neonatal, adult and aged stages. A majority of single cells were largely clustered by organs at E16.5 and P5, but intermixed at adult and aged stages (Fig. 2A). We calculated the organ-specific signature score to examine the epigenomic memory of each organ with time. Profound organ-specific features determined by single-cell H3K27ac signals were observed in the developmental lives (E16.5 and P5) (Fig. 2B). Notably, robust H3K27ac signals were evident around TF genes, thus likely reporting tissue developmental origins of brain (*Olig2, Otx1, Neurog1*)(*60-62*), heart (*Tbx5, Mef2c, Tbx20*)(*63-65*), kidney (*Hoxa9, Hoxa11, Pax2*)(*66-68*), liver (*Hnf4a, Foxa2, Hnf1a*)(*69, 70*), and lung (*Foxf1, Sox9, Foxp2*)(*71-73*) (Fig. 2, C to G, and fig. S3, A to E). In addition, to analyze the epigenomic heterogeneity of EC lineage cells within stages, we identified different clusters for organs in each stage (fig. S4A), and performed correlation analysis between clusters as well as Jaccard similarity for single cells. In line with the aforementioned, the aged stage showed the lowest similarity across and within clusters and single cells, indicating a dramatic epigenetic change during ageing (fig. S4, B and C).

**Fig 2.**
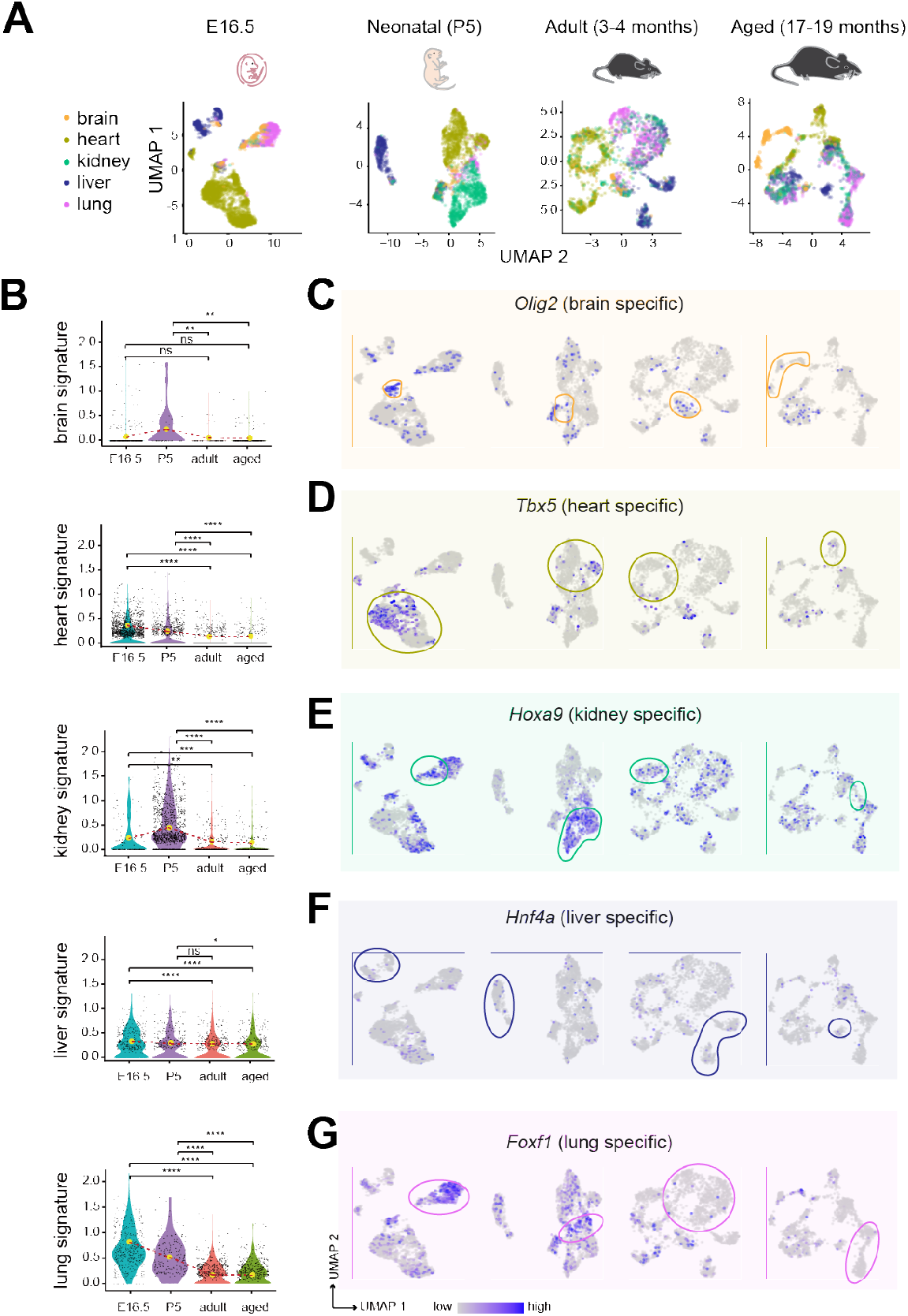
Epigenomic memory of dynamic endothelial lineages at developmental transcription factors reflects their origin of organ. **(A)** UMAP visualization of single cells colored by organs at different stages. Cell number: n=3,322 in E16.5, n=3,062 in P5, n=2,224 in adult, and n=1,919 in aged stage. **(B)** Violin plots showing the organ developmental TF signature score at different stages. One-tailed t test was used to determine the statistical significance, ns, not significant, *\*P <* 0.05, *\*\*P* < 0.01, *\*\*\*P* < 0.001, and *\*\*\*\*P <* 0.0001. The average score of each stage was linked by red dashed line. The H3K27ac signals were more profound at E16.5 or P5 but later diminished to a varying degree in different organs, indicating that the epigenomic memory at organ-developmental TF genes are temporally and differentially regulated across organs. **(C** to **G)** UMAP showing H3K27ac signals around organ-developmental TF genes. Cells with the organ specific TF signature are outlined, from brain (C), heart (D), kidney (E), liver (F) and lung (G).

To explore the contribution of different EC lineage subtypes to tissue specialized functions for development and homeostasis, we clustered single-cell H3K27ac CoBATCH data for each organ across stages. Cell identities were annotated based on several known marker genes together with the previously established gene signatures (table S5) for organ function (Fig. 3, A and B, and fig. S5A). For example, we identified a cluster corresponding to blood–brain barrier (BBB) ECs shielding the brain from circulating blood components(*74*), annotated by H3K27ac signals around BBB related genes *Slc2a1*(*75*) and *Foxq1*(*76*). Diverse EC and non-EC subtypes in heart were also identified(*77*). Notably, a proangiogenic EC subpopulation was linked to the signaling of VEGF, primarily facilitating the growth of blood vessels(*78*). *Cgnl1* was known as a unique marker for endocardial cells, phenotypically distinguished from vascular ECs(*32*). *Dach1* as a known marker in regulating coronary vasculature growth by potentiating arterialization, facilitated the definition of the function specialization in heart(*35*). Further, fenestrated ECs, specialized for their unique role for filtration across the glomerular capillary wall in kidney, were characterized by *Plvap(79*) and *Plxnb1*(*80*). Liver sinusoidal endothelial cells (LSECs) form a unique barrier and are involved in metabolic maintenance and organ homeostasis(*81*). These highly specialized ECs were identified by highest H3K27ac signals around the *Stab1*(*82*) and *Id1*(*83*) gene loci. We also defined pulmonary alveolar type I (AT1)-like cells in lung by typical markers (*Hopx*(*84*) and *Ager*(*85*)), which are specialized for efficient gas exchange (Fig. 3C and fig. S5B). The identity of each cell subtype with specialized functions was also examined alongside projection of the pan-EC signature score and specific inspection of H3K27ac signals at *Pecam1* and *Cdh5* (fig. S5C).

**Fig 3.**
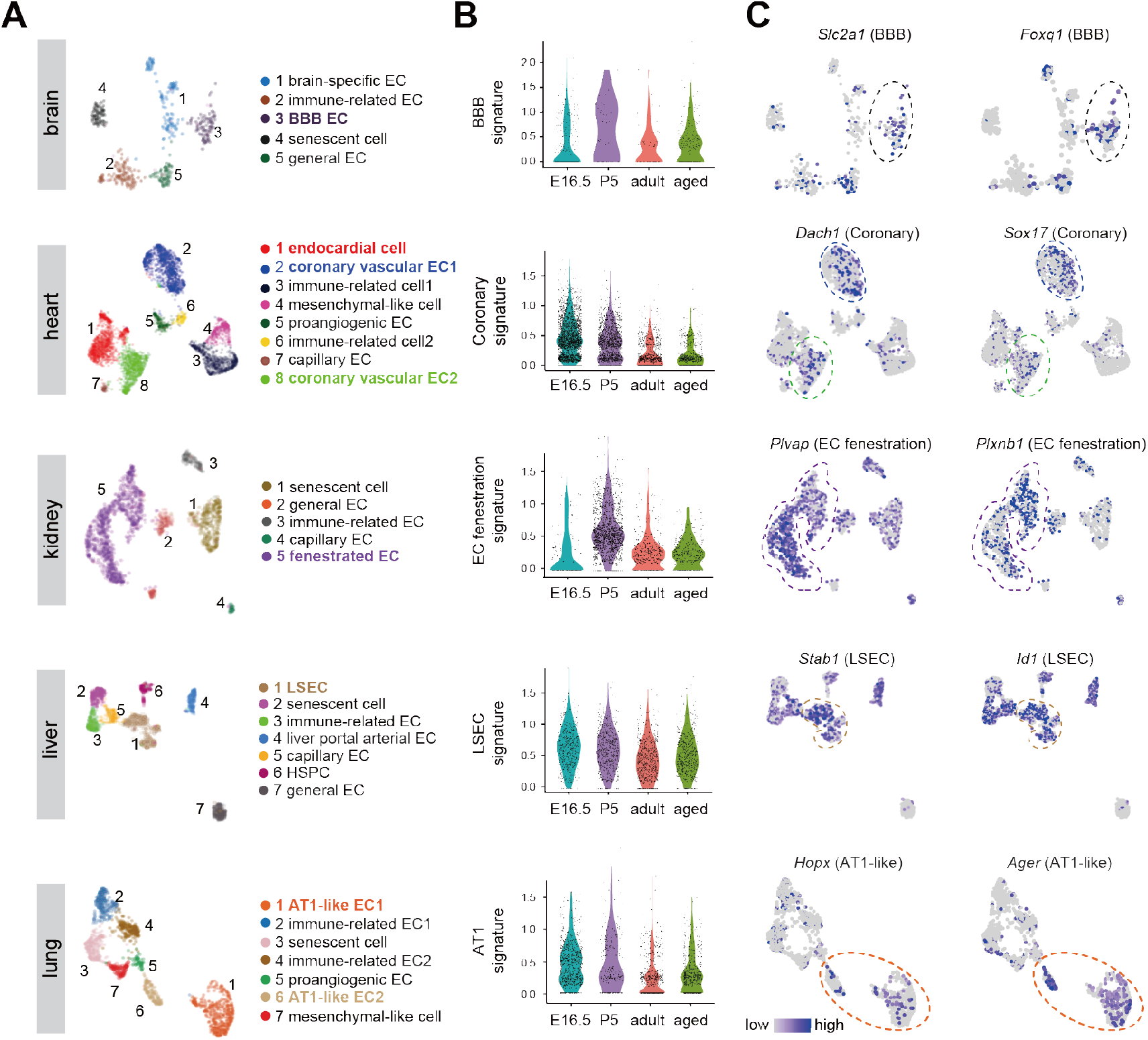
Epigenomic memory of dynamic endothelial lineages nominates organ specialized functions. **(A)** UMAP visualization of EC lineage cells from H3K27ac CoBATCH data colored by cell types within each organ. The organ-specific ECs with functional specialization were annotated and highlighted by multicolor in (A), n = 460, 4,774, 1,733, 2,302, and 1,692 cells for brain, heart, kidney, liver, and lung, respectively. EC, endothelial cells; BBB, blood-brain barrier; LSEC, liver sinusoidal endothelial cell;AT1, pulmonary alveolar type I. **(B)** Violin plots showing the signature score related to the specific organ function at different stages. **(C)** Feature plots of H3K27ac signals around organ-specific function genes in the subtypes. Cells with the organ specific TF signature are outlined, from brain (*Slc2a1, Foxql*), heart (*Dachl, Sox17*), kidney (*Plvap, Plxnbl*), liver (*Stabl, Id1*), and lung (*Hopx, Ager*).

### Identification of temporally regulated transcription factors implicated in EC fate change

To recapitulate the developmental plasticity of the EC fate within organs, we reconstructed pseudotime trajectories for each organ based on single-cell H3K27ac profiles using Monocle 3(*86*). The predicted pseudotime was in good agreement with the real developmental stages within each organ (Fig. 4A and fig. S3A). We next sought to investigate the dynamic change in TF motifs along pseudotime. Interestingly, we observed dynamic patterns for organ-specific TFs (HES1 and NR2E1 in brain, ISL1 and NKX2-5 in heart, PAX2 and HOXA11 in kidney, HNF4A and GATA4 in liver, and FOXD8 and HOXF1 in lung), correlated with corresponding organ development (Fig. 4B). Further, the oscillating ERG and TWIST1 motif scores along pseudotime exhibited a reverse correlation within each organ (Fig. 4C). This partly validated the pseudotime ordering of single cells and also supported the important role of EndMT during organogenesis. Unsupervised k-means clustering was able to nominate a number of temporally dynamic TFs within distinct regulatory regions along pseudotime (Fig. 4D). Together, these results revealed the temporally regulated TF activity closely associated with the EC fate transition, including EndMT.

**Fig 4.**
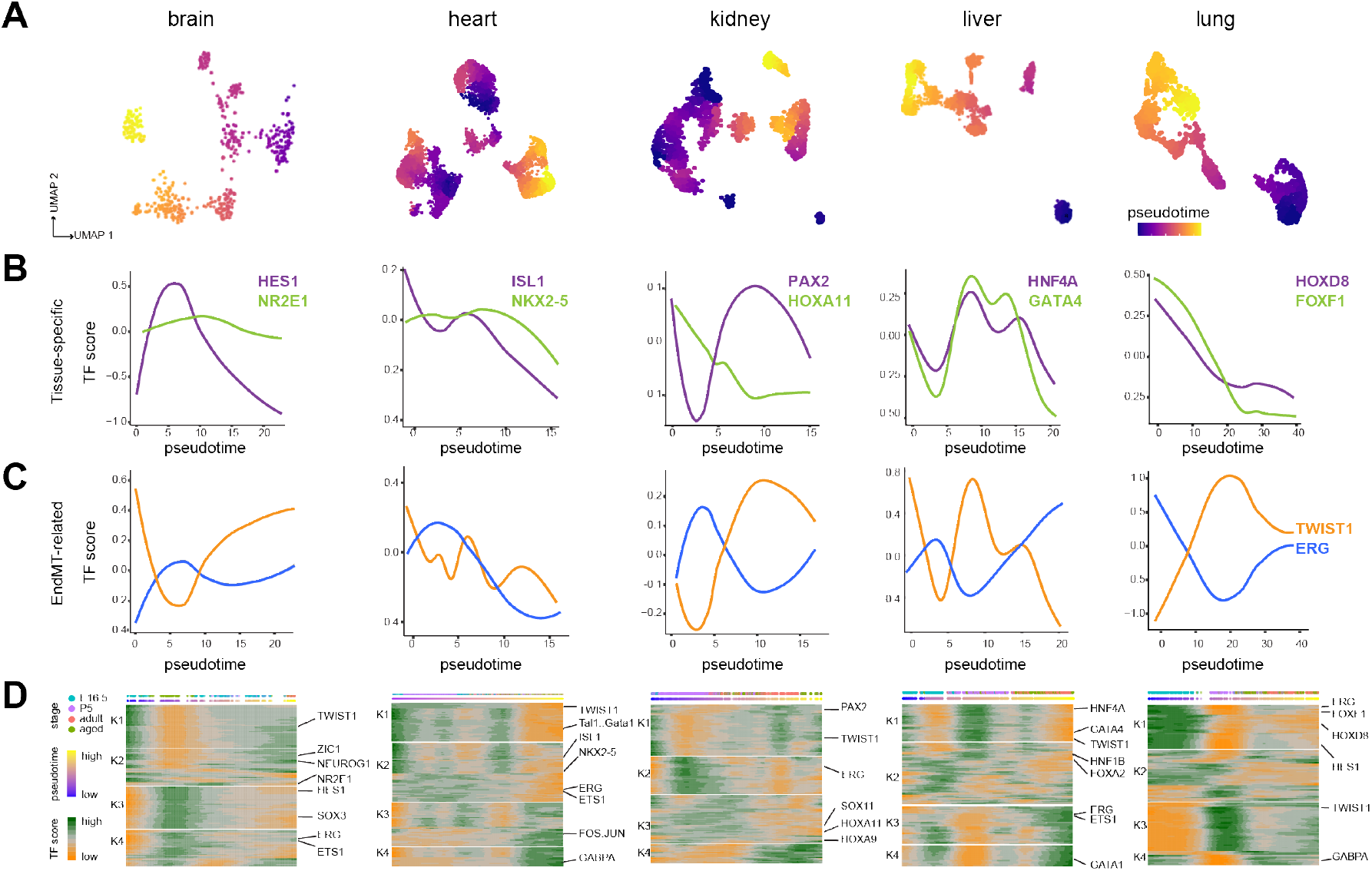
Identification of temporally regulated organ-specific transcription factors implicated in endothelial cell fate transition. **(A)** Pseudotime trajectories of single cells in each organ using LSI-based 50 dimensionalities of H3K27ac profiles by Monocle 3. **(B)** The dynamics of motif scores of known organ-specific TFs along pseudotime identified in (A). Representative TFs shown here include HES1 and NR2E1 (brain), ISL1 and NKX2-5 (heart), PAX2 and HOXA11 (kidney), HNF4A and GATA4 (liver) and HOXD8 and FOXF1 (lung), exhibiting a similar pattern of dynamics along pseudotime. TF motif score which represents the motif activity was calculated as the motif deviations in H3K27ac signals across the peaks using chromVAR. **(C)** The dynamics of motif scores of the EndMT related TFs ERG and TWIST1 along pseudotime identified in (A). The dynamics of the EC-specific ERG and mesenchymal-specific TWIST1 along pseudotime was reversely correlated. EndMT, endothelial-to-mesenchymal transition. **(D)** Heatmap showing k-means clustering of smoothed scores of TF motifs (JASPAR2020 database) along pseudotime in each organ. Dots on the top two rows represent single cells. The colors in the first row show real age information, and the colors in the second row indicate pseudotime.

### EndMT process reinforced by bivalent chromatin co-binding of EC- and MC-enriched TFs

EndMT is a cell fate dynamic process involved in many biological processes, such as heart valve formation, tissue regeneration and ageing(*87, 88*).we next sought to perform ERG and TWIST1 CoBATCH profiles to explore the single-cell TF binding landscape during EndMT. Similarly, hearts from Cdh5^CreERT2^::Rosa26^tdTomato/+^mice induced by tamoxifen at P1 and harvested at P5 for FACS sorting for tdTomato^+^ cells (Fig. 5A). After stringent filtering to remove a significant portion of cells expressing neither ERG nor TWIST1, 457 and 329 single cells were retained for subsequent analyses. We performed LSI and UMAP with combined datasets from independent ERG and TWIST1 CoBATCH experiments and identified two major subpopulations including ECs and MCs defined as non-ECs (Fig. 5, B and C). No evident batch effect was observed when clustering all single cells from different datasets (Fig. 5, B and C). We also confirmed bona fide peaks from aggregate ECs and MCs, and characterized their potential functions (fig. S6, A to C). To determine the differential TF motif activity from ERG and TWIST1 binding sites during EndMT, we used chromVAR to calculate motif scores(*89*). Expectedly, the ERG/ETS1 motif was highly enriched in ECs, and the high motif scores of ZEB1(*90*), SNAI1, SNAI2(*87*) were observed in MCs (Fig. 5, D and E). Interestingly, beyond this, there was also the high ERG motif score in the EC cluster identified from TWIST1 CoBATCH profiles, and vice versa (Fig. 5E). This finding suggested that ERG was likely recruited to a subset of TWIST1 peaks via indirect mechanisms and so was TWIST1.

**Fig 5.**
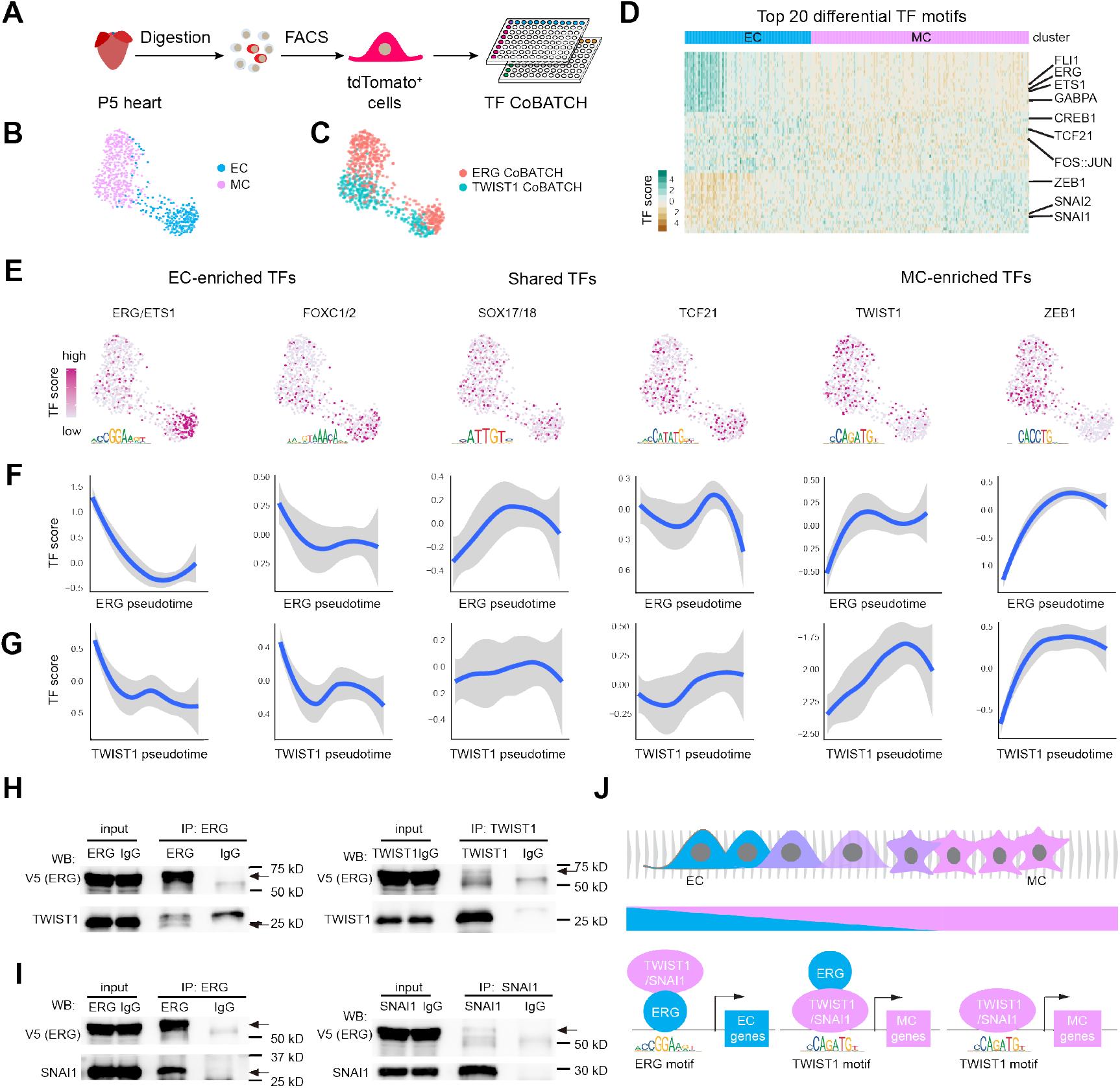
Single cell TF CoBATCH reveals the EndMT process reinforced by bivalent chromatin co-binding of EC- and MC-enriched TFs. **(A)** Experimental design of TF (ERG and TWIST1) CoBATCH experiments in P5 heart from Cdh5^CreERT2^::Rosa26^tdTomato/+^rnice. **(B** and **C)** UMAP visualization of single cells from ERG and TWIST1 CoBATCH data. Single cells were colored by cell types (B) and CoBATCH experiments (C). n = 457 and 329 cells for ERG and TWIST1 CoBATCH data, respectively. **(D)** Heatmap showing TF scores enriched in each subpopulation identified in (B). Top 20 TFs ranked by P value were shown here. **(E)** Feature plots showing the TF motif enrichment scores for representative EC (left), shared (middle), MC (right) TFs. EC, endothelial cell; MC, mesenchymal cell. **(F** and **G)** The dynamic changes of TF scores for representative factors in (E) along ERG (F) and TWIST1 (G) pseudotime during EndMT. The blue lines are the smoothed conditional means and the grey shadow indicates the 95% confidence interval. **(H)** Co-immunoprecipitation showing ERG and TWIST1 interaction. Immunoprecipitation with antibodies against ERG (left) or TWIST1 (right) in HEK293T cells transfected with TWIST1 and V5 tagged ERG and Western blotting with indicated antibodies V5 or TWIST 1. Arrows indicate the full-length V5-ERG (top) and TWIST 1 (bottom), respectively. **(I)** Co-immunoprecipitation showing ERG and SNAI1 interaction. Immunoprecipitation with antibodies against ERG (left) or SNAI1 (right) in HEK293T cells transfected with SNAI1 and V5 tagged ERG and Western blotting with indicated antibodies V5 or SNAI1. Arrows indicate the full-length V5-ERG (top) and SNAI1 (bottom), respectively. **(J)** Working model summarizing transient EC- and MC-TFs interaction mediated bivalent chromatin co-binding during the process of EndMT. Shown are the ERG and TWIST1 motif, representing the ETS TF family and MC TFs, respectively.

Next, to gain in-depth understanding of the mechanism, we carried out pseudotime analysis and revealed a temporally progressive change in the motif score of EC-enriched TFs (ERG/ETS1 and FOXC1/2), shared (SOX17/18 and TCF21), and MC-enriched TFs (TWIST1 and ZEB1). This yielded consistent kinetics for the cell-type specific TFs along pseudotime between ERG and TWIST1 profiles (Fig. 5, F and G). Strikingly, a significant fraction of single cells from ERG CoBATCH profiles presented the high TWIST1 but lowest ERG motif scores, suggesting that we captured chromatin occupancy of TWIST1 via ERG tethering within these cells. Likewise, this was also the case with identified ERG binding in TWIST1 CoBATCH profiles. Collectively, these results lead to a hypothesis that the physical protein interaction between EC and MC TFs, such as ERG/TWIST1, may give rise to the bivalent chromatin co-binding at single-cell resolution.

To validate such protein–protein interaction, we performed reciprocal co-immunoprecipitation assays for ERG/TWIST1 and ERG/SNAI1 (another MC-enriched TF). Indeed, our results showed that both TWIST1 and SNAI1 immunoprecipitation in HEK293T cells co-purified ERG proteins, and vice versa, confirming the interactions (Fig. 5, H and I). Taken together, these data support a model that the TF complex bind to the regulatory elements of EC genes via the ERG binding activity at the onset of EndMT while with progression to those of MC genes via the TWIST1/SNAI1 binding activity (Fig. 5J).

### H3K27ac and H3K36me3 CoBATCH profiles distinctly report endothelial lineage senescence

Accumulating evidence shows that the ageing process is associated with epigenetic alterations within each organ across the lifespan(*91*). To investigate the coupling between the temporal dynamic landscape of different histone modifications and ageing features, we first analyzed H3K27ac and H3K36me3 (largely deposited at actively transcribed gene body(*92, 93*)). CoBATCH profiles in hearts from development through aged stages. A two-dimensional representation of H3K27ac profiles revealed a prominent separation of EC lineage cells into early (E16.5), middle (P5) and late (adult, aged) phases (Fig. 6A). We next asked how cellular composition changed during the ageing process. At the earlier time point (E16.5, P5) in H3K27ac profiles, the cell population was mainly composed of cardiac ECs (95% and 92%). Starting from P5, immune-related cells were present with a marked increase at adult and aged stages (Fig. 6B). The enhanced immune response with age was also witnessed by analyzing bulk RNA-seq data from different ages of adult hearts (fig. S7A). Although a similar trend in cellular composition was observed in H3K36me3 CoBATCH profiles, these effects in cardiac EC and immune-related cell composition were more pronounced in H3K27ac profiles (Fig. 6B). To further characterize the dynamics during endothelial-related biological processes mounted in ageing, we calculated the H3K27ac and H3K36me3 values for pan-EC, artery, capillary, venous, lymphatic, angiogenesis, EndMT, immune-related, and housekeeping gene signatures. Expectedly, pan-EC signatures showed a steady decrease while immune-related and EndMT signatures manifested an evident increase along pseudotime within H3K27ac profiles in each organ (Fig. 6C and fig. S7B). However, this finding was not observed in H3K36me3 profiles (Fig. 6C). This phenomenon was further supported by the fact that single cells from H3K27ac profiles along the descending EC score were accompanied by an increment in the immune-related signature score during ageing. Again, this was absent with H3K36me3 profiles (Fig. 6D).

**Fig 6.**
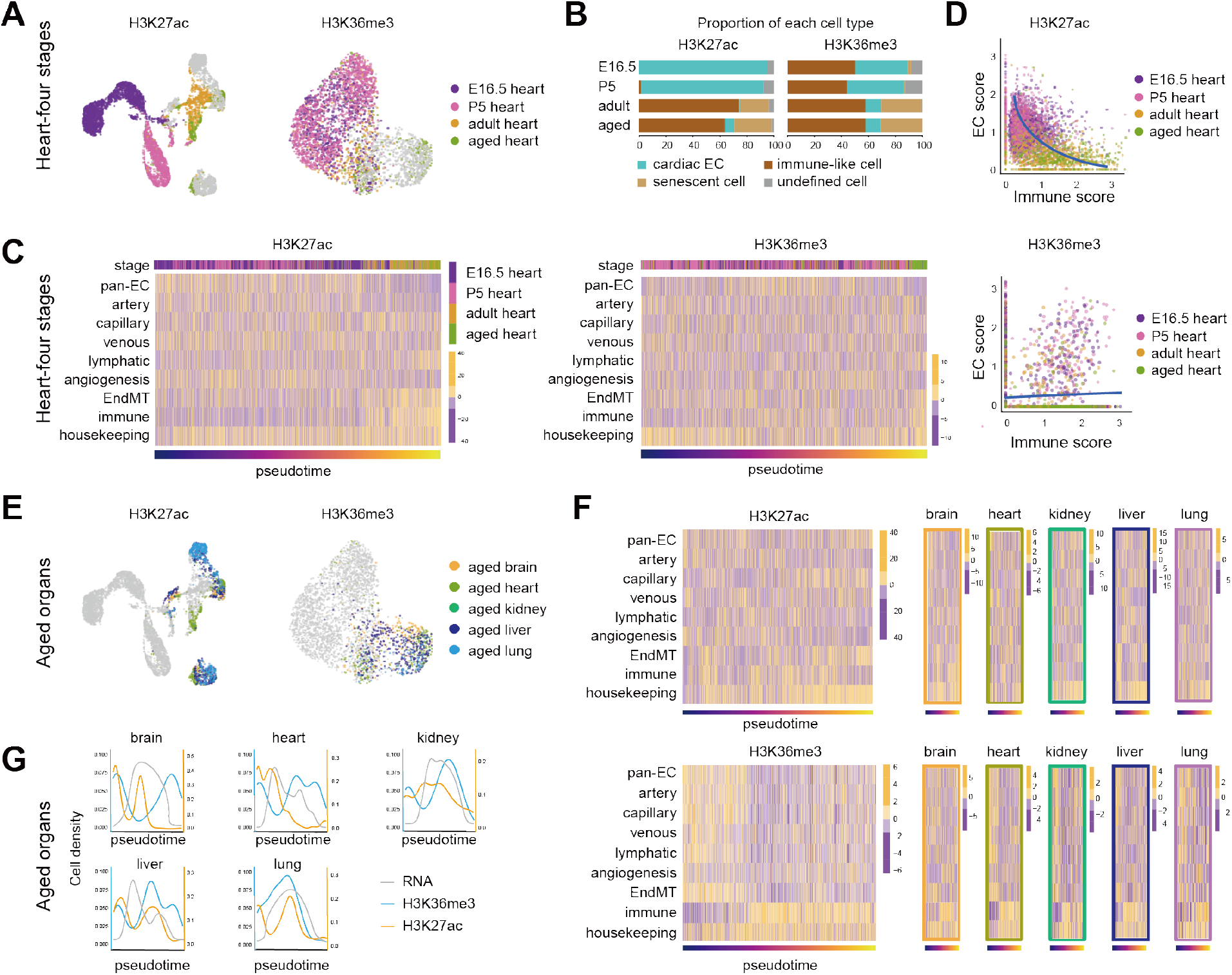
H3K27ac and H3K36me3 profiles disparately report endothelial lineage senescence. **(A)** UMAP visualization of EC lineage cells from H3K27ac (left) and H3K36me3 (right) CoBATCH data. Single cells from heart were colored by four stages, and others in grey are from aged brain, kidney, liver, and lung. **(B)** Proportion of each cell type for H3K27ac (left) and H3K36me3 (right) profiles in four stages of hearts. **(C)** Heatmaps showing single cell H3K27ac (left) or H3K36me3 (right) signals at peaks around nine signature module genes in heart ECs. Modules include pan-EC, artery, capillary, venous, lymphatic, angiogenesis, EndMT, and immune and housekeeping genes. Top row showing real stage annotation and bottom row showing pseudotime ordering. **(D)** Scatter plots showing EC (y-axis) and immune score (x-axis) calculated using H3K27ac (top) and H3K36me3 (bottom) profiles for single cells in heart colored by stages. **(E)** UMAP visualization of EC lineage cells from H3K27ac (left) and H3K36me3 (right) profiles. Single cells in the aged stage were colored by organs, and others in grey are from the stages of E16.5, P5, and adult. **(F)** Heatmaps showing single cell H3K27ac (top) or H3K36me3 (bottom) signals at peaks around nine signature module genes in ECs from aged organs. **(G)** The cell density distribution from H3K27ac (blue line), H3K36me3 (orange line) CoBATCH and RNA profiles (grey line) along pseudotime in each organ. The scRNA-seq data of five organs for 18-month mice used here were downloaded from the Tabula Muris Senis database.

Next, we focused on the aged stage across five organs (brain, heart, kidney, liver, and lung) to examine the short-range epigenetic change compared to the long-range window analyzed above (Fig. 6E). Strikingly, by calculating the signature scores, we observed the more profound transition in H3K36me3 profiles compared to H3K27ac, suggesting that single-cell H3K36me3 exhibited a shorter epigenomic memory than H3K27ac during ageing (Fig. 6F). To distinguish the transitioning modes, we also uncovered that a significant fraction of cells was distributed at the nearly ending time (more senescent) in H3K36me3 profiles in all organs except for lung. However, a majority of cells were found at the early phase along pseudotime of senescence in H3K27ac profiles within each organ (Fig. 6G). As a control, we also performed the similar analysis with single-cell transcriptomic data from the Tabula Muris Senis(*14*), finding a more continuous distribution of cells (fig. S7, C to F). Together, our data support that the EC lineage senescence experiences a gradient increase in the immune response determined by single-cell H3K27ac profiles in the long-range time window. Nonetheless, H3K36me3 regulation is more sensitive in this regard in the short period of time.

## Discussion

We reconstructed the single-cell epigenomic trajectories of plastic EC lineages in the five representative mouse organs across developmental and ageing stages by CoBATCH profiling of histone modifications and transcription factors. Our study differs from other recent single-cell atlas researches in several aspects. A vast majority of recent single-cell reports leveraged scRNA-seq for cataloging cell types from specific tissues or across many organs in adult or embryonic stages(*5, 94-98*). Other studies capitalized on variant single-cell ATAC methods to profile chromatin accessibility for understanding of key regulatory elements in transcriptional regulation and cell differentiation in snapshot data from mouse or human tissues(*21, 22*). More recently, an elegant study specifically interrogated the heterogeneity of ECs across mouse adult tissues, identifying 78 EC subtypes defined by transcriptome across different vascular beds(*15*). Aside from distinct layers of molecular information detected, our study explored the real EC lineage dynamic trajectories over 4-day lineage tracing in several stages from development through adult to aged stages. Given the reported high plasticity of ECs *in vivo*, the rational is that 4-day tracing rather than single snapshot of cells actively expressing EC markers would offer better opportunities to decipher the EC developmental origin and epigenetic basis of cell fate regulation. Importantly, we examined not only EC lineages in organogenesis but also their senescence process. Finally, cross-tissue comparisons of these broadly distributed EC lineages yielded distinctive epigenomic features in cell fate plasticity and further remodeling.

Our work demonstrates the power of CoBATCH to illustrate not only the epigenetic heterogeneity and plasticity of ECs in mouse organs across the lifetime, but also the mechanisms that underpin cell fate transition. For example, directly assaying TF chromatin occupancy by CoBATCH, rather than computationally inferring putative TF motifs within regulatory regions(*99*), revealed that the EndMT process was reinforced by bivalent chromatin co-binding of otherwise mutually exclusive EC- and MC-TFs. Further, single-cell temporal trajectories of H3K27ac and H3K36me3 profiles enabled faithfully capturing the finetuning EC lineage senescence in a long- and short-time window.

New insights were also gained through epigenomic tracing of developmental origin of EC sublineages. One major goal in developmental biology is to trace a cell back to its origin. Despite a large body of studies have demonstrated that many tissue-resident ECs are of mesodermal origin, different origins of ECs in several tissues have been reported(*100-105*). For example, mesoderm derived vessel plexus contributes to hepatic ECs(*101*); endocardial cells in the developing heart contribute to part of liver vasculature, indicating a potential cross-organ regulation through direct vascular contribution during organogenesis(*102*). Our data uncovered developmental origin of ECs based on epigenomic memory, but also witnessing flexible time lengths across organs. For instance, early EC lineages retained the strong H3K27ac memory around cognate pioneer TFs expressed in tissue progenitor cells, such as *Hoxa9* in kidney, *Hnf4a* in liver, and *Foxf1* in lung. As a caveat, these inferring results need future investigation through genetic lineage tracing.

Looking forward, further expanding datasets by including more EC lineage cells covering the entirety of mouse organs and adding epigenomic layers across many other histone modifications would undoubtedly improve exploration of rarer but regenerative EC or its derived non-EC subtypes. We foresee that will reveal a full organism-wide view of the epigenetic landscape of the widely distributed EC lineage plasticity. This will bring us closer to produce organotypic ECs with therapeutic applications. Furthermore, concurrent measurement of epigenome and transcriptome in single cells would greatly deepen understanding of the complex link between multilayered molecular information and EC differentiation, remodeling and plasticity in development and diseases.

## Acknowledgments

We thank all members of the He lab for critical comments on this manuscript. Part of the analyses was performed on the High Performance Computing Platform of the Center for Life Sciences, Peking University. We thank the flow cytometry Core at National Center for Protein Sciences at Peking University, particularly Liying Du and Jia Luo, for technical help.

## Author contributions

A.H. conceived and designed the study. X.Y. designed and performed all experiments. Y.L. and H.X. performed the computational analyses, supervised by A.H. X.L. provided computational support. H.X., Y.L. and A.H. wrote the paper with input from all other authors. All authors participated in data discussion and interpretation.

## Supplementary Materials

Materials and Methods

Figs. S1 to S7

Tables S1 to S5

